# Characterizing the epigenetic landscape of cellular populations from bulk and single-cell ATAC-seq information

**DOI:** 10.1101/567669

**Authors:** Mariano I. Gabitto, Anders Rasmussen, Orly Wapinski, Kathryn Allaway, Nicholas Carriero, Gordon J. Fishell, Richard Bonneau

## Abstract

Given its ability to map chromatin accessibility with single base pair resolution, ATAC-seq has become a leading technology to probe the epigenomic landscape of single and aggregated cells. Understanding ATAC-seq data presents distinct analysis challenges, compared to RNA-seq technologies, because of the relative sparseness of the data produced and the interaction of complex noise with multiple chromatin structure scales. Methods commonly used to analyze chromatin accessibility datasets are adapted from algorithms designed to process different experimental technologies, disregarding the statistical and biological differences intrinsic to the ATAC-seq technology. Here, we present a Bayesian statistical approach, termed ChromA, to analyze ATAC-seq data. ChromA annotates the cellular epigenetic landscape by integrating information from replicates, producing a consensus de-noised annotation of chromatin accessibility. Our method can analyze single cell ATAC-seq data, improving cell type identification and correcting many of the biases generated by the sparse sampling inherent in single cell technologies. We validate ChromA on several biological systems, including mouse and human immune cells and find it effective at recovering accessible chromatin, establishing ChromA as a top preforming general platform for mapping the chromatin landscape in different cellular populations from diverse experimental designs.

## Main

The genome of eukaryotic cells is tightly packed into chromatin [1] with only a fraction of chromosomal regions accessible within any given cell population at any given time. Chromosomal accessibility plays a central role in several nuclear processes including the regulation of gene expression and the structure and organization of the nucleus [2]. Transcription factor proteins (TFs) bind to accessible DNA regions to control the expression of genes [3] and to inaccessible chromatin, altering the accessibility of targeted regions [4]. Differential expression and regulation of TFs act as a combinatorial code that gives rise to the wide repertoire of cellular phenotypes observed in mammalian organisms [5], [6].

The heterogeneity of cellular types and the importance of regulation of chromatin structure is well illustrated in the vertebrate immune system. The immune system is composed of well-defined cellular types for which the action of TFs and their targets are known for a subset of key transcriptional regulatory networks [7]. Master regulators dictating the diversification of different lineages of T lymphocytes in the immune system have been extensively characterized [8],[9],[10] and recently, multiple new regulators for the RORg TF expressing lineage, Th17, have been identified [12],[13]. This functional and mechanistic context combined with the extensive experimental data available for these cell types, make them an ideal platform to quantify accessible chromatin and associated regulatory elements.

The development of high-throughput chromatin accessibility assays (ATAC-seq) has enabled the analysis of chromatin accessible regions, the discovery of nucleosome positions and the characterization of transcription factor occupancy with almost single base pair resolution [14]. In part due to the small initial starting material (on the order of 10000 cells) and from a desire to query the chromatin structure of particular cellular types, ATAC-seq has become widely adopted. Recent advances have improved the technique and enabled the mapping of the accessible chromatin landscape of individual cells [15]. This, in turn, raises the possibility of both describing the variability of chromatin accessibility and enabling classification of cellular types based on their chromatin structure [16] [17].

Compared to techniques for assaying RNA expression at single-cell and bulk levels [18], chromatin profiling presents considerable challenges. Specifically, there is at present no systematic approach for characterizing the variability of read counts present at each base within an accessible region. Although many genes localize in already known genomic loci, such as annotated exons and introns, several additional genomic elements remain uncharacterized and much of the chromatin landscape in many cell-types exhibits an unknown structure [19]. This unstructured landscape combined with the extreme sparsity inherent in the assay (caused by the fact that the maximum information available at each genomic locus is at most two reads per cell), inherently limits the signal-to-noise ratio when assaying chromatin accessibility. The correct annotation of the chromatin landscape is of paramount importance in the identification of different cellular types and the linkage of distal elements to promoter regions. A fact supported by studies that identified distal enhancer elements as regulatory regions driving lineage commitment [20], [21]. Thus far, ATAC-seq experiments have primarily been analyzed using statistical tools developed to understand MNase or DNAse assays [22] [23]. As a consecuence, ATAC-seq lacks a proper statistical model suited to its analysis. Such a model should consider not only a data-driven read generative model but also the high degree of biological variability in the size of accessible regions, which varies from a few tens of base pairs to thousands.

Here, we present ChromA, a Bayesian statistical approach that models ATAC-seq information to infer chromatin accessibility landscape and annotate open (accessible) and closed (inaccessible) chromatin regions. ChromA harnesses recent developments in hidden semi-Markov models to create a scalable statistical inference method that can be applied to genome wide experiments [24]. When modeling experimental replicates, ChromA is able to integrate information from different experiments and create a consensus chromatin annotation. To validate our method, we used Th17 bulk [12], A20 and GB12878 single-cell datasets identifying accessible chromatin and its regulatory elements, establishing ChromA as an effective platform for mapping the chromatin landscape in different cellular populations.

## Results

### A Hidden Semi-Markov Model for Chromatin Accessibility Annotation

ChromA (Chromatin Accessibility Tool) is a probabilistic graphical model developed to annotate chromatin regions as open (accessible) or closed (not accessible) when experiments are performed on pooled (bulk), single cells or a combination of both. Our algorithm takes as an input ATAC-seq aligned sequencing reads (.bam files) or locations of Tn5 binding events (.tsv files) and recovers chromatin accessibility annotations and quality control metrics for the dataset (Figure 1A).

**Figure 1:**
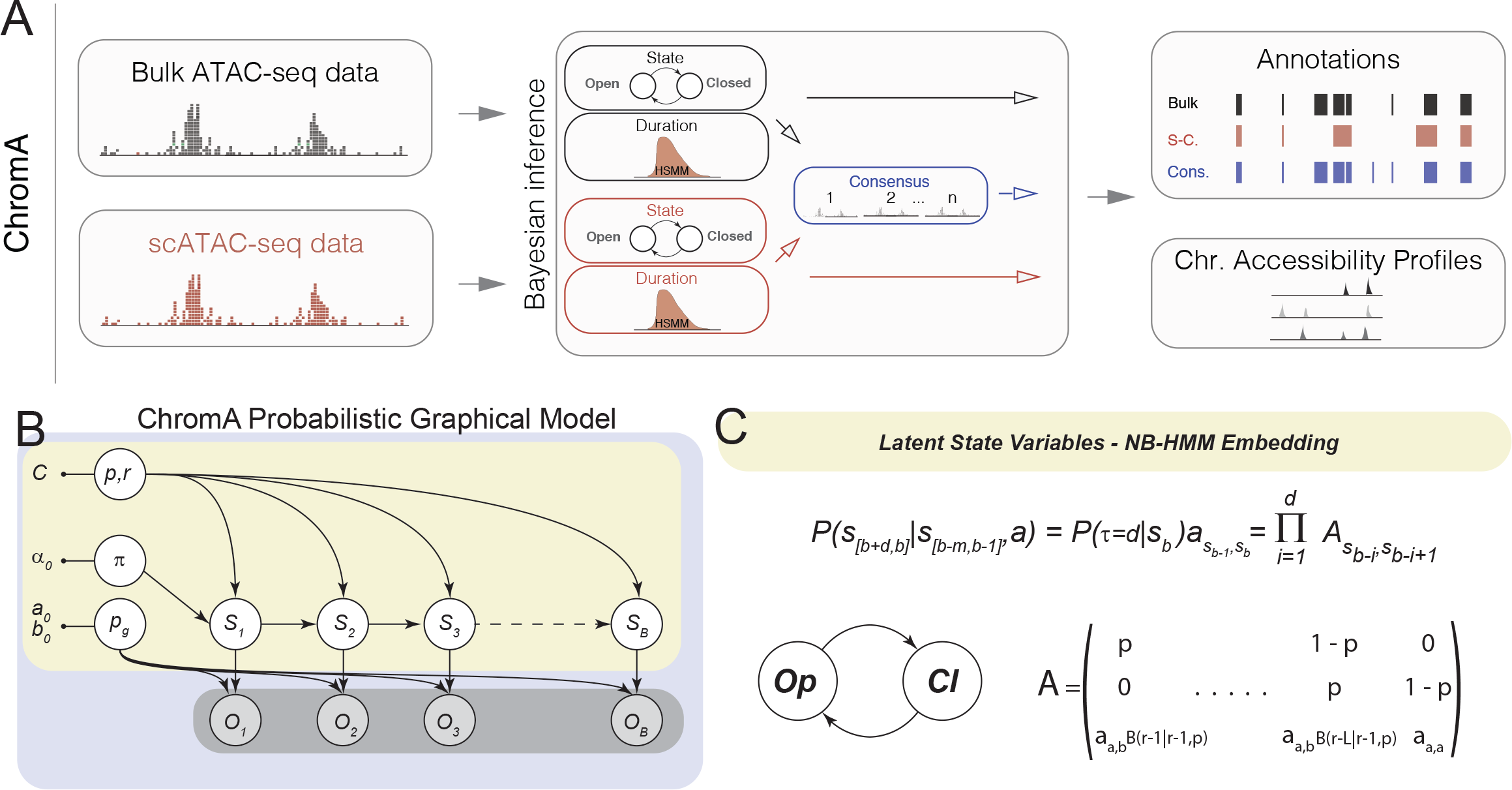
Overview of Chromatin Accessibility Annotation Algorithm. (a) ChromA is an easy to use algorithm that combines single and multiple bam files (raw reads) or tsv files (list of Tn5 binding events) to create chromatin accessibility annotations. ChromA produces different metrics to inform users the quality of the processed dataset. (b) Probabilistic graphical model describing ChromA’s structure. In this representation, nodes describe random variables and arrows depict dependencies among the variables. ChromA models the number of Tn5 binding events observed at each base using observed variables *X* (representing the number of binding event), and latent variables *S* (representing chromatin state). Subscripts denote base position, ranging from 1 to the length of a chromosome, *B*. Observed variables *X* are modelled using a geometric distribution with parameter *p_g_*. Chromatin state variables *S* are subjected to semi-Markovian dynamics, depending on the previous chromatin state. *π*described the initial chromatin state, and p and r characterize the semi-Markovian transition matrix dictating chromatin state context. (c) Our ATAC-seq pipeline using bulk measurements annotates chromatin using two states, open Op, or closed Cl. Both states are characterized by semi-Markovian dynamics. The probability of annotating chromatin in bases *b* to *b + d* given previous chromatin states depends on two factors: the probability of transitioning between states, symbolized by transition matrix *a*, and the probability of dwelling in the new states during *d* bases. When the duration is characterized by a negative binomial (NB) distribution with parameters p and r, the transition matrix can be re-written using an embedding matrix *A*. In the figure, we reproduce a simple transition matrix *A* in which *p* and *r* are the NB parameters, *a* is the transition matrix between states, and *B* represents the binomial coefficient.

ChromA is based on a Bayesian statistical model that encompasses a set of latent variables (*S*) representing accessibility at each base and a set of observations (*O*) composed by the reads (Figure 1B). In our model, the chromatin state of each base is a binary variable representing two chromatin conditions, open (*S_b_ = 1*) and closed (*S_b_ = 0*). Bayesian inference creates posterior estimates of model’s parameters by combining our prior belief about parameter values with the likelihood of the observations being generated by the model. In our case, ChromA aims to estimate posterior chromatin state by combining our prior belief on the accessibility of each base with the likelihood of generating the observed reads.

To model the duration of accessible regions from ATAC-seq experiments, we reason that contextual information plays a key role in defining each base’s annotation. Hidden Markov models (HMM) enjoy a huge popularity in genomic applications because they integrate contextual information into their statistical postulates by adding dependencies between adjacent base pairs. This contextual dependency translates into a modelling assumption about the duration of each state, *d*, given in HMMs by a geometric distribution [25] (eqn. 1).

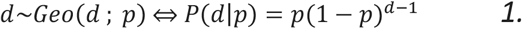

where the symbol ∼ denotes that duration variable *d* is distributed according to a geometric distribution and *p* is the probability of dwelling in the state before transitioning away into another state. To improve upon the duration behavior of standard HMMs we propose to model the duration of each accessible region through a hidden semi-Markov model (HSMM) that exhibits a negative binomial (NB) duration distribution [26] (eqn 2).

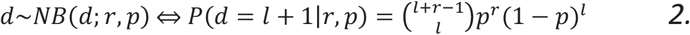

The NB distribution has two parameters: an integer parameter *r* > 0, and a probability parameter 0<*p*<1. We use this distribution to capture the notion that chromatin accessible regions might require a certain characteristic length to host the cis-regulatory transcriptional machinery necessary for accessing DNA-binding domains. The maximum or mode of a negative binomial distribution is given by its parameters 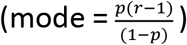. This is contrary to models based on the geometric distribution for which the maximum is fixed and always reached at 1 (supplementary figure 1a). An example of better fitness when the NB distribution is used in genomic applications is given by the characteristic length of a gene. The negative binomial distribution can be used to effectively describe gene lengths across different chromosomes, consistent with the idea that genes might need a certain characteristic length to create functional proteins (supplementary figure 1b).

Recent developments in approximate posterior calculation provide efficient techniques for the estimation of HSMM parameters. These techniques are advantageous when the duration of HSMM states are distributed according to a NB distribution [24]. To harness the advantage of such developments, we focus on the parameter that encodes the duration of each state in HMMs and HSMMs, the transition matrix. An HMM possess a simple transition matrix, *A_ij_*, denoting that the probability of transitioning into a new state *j* at base b depends on the state *i* of the previous base (Markovian Property) (eqn. 3).

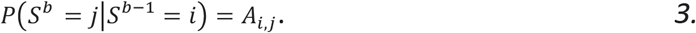

The transition matrix of a HSMM, *A′*_i,j_, under the assumption of independence on the previous state duration, can be written using two terms: the probability of transitioning into a new state, E_F,G_, and the probability of dwelling in the new state for a duration of d bases, *P*(τ = *d*|*S^t^* = *j*) [eqn. 4].

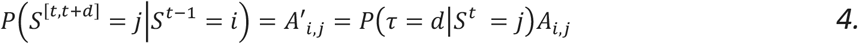

To facilitate inference, we begin by re-writing the NB distribution as a sum of shifted geometric distributions.

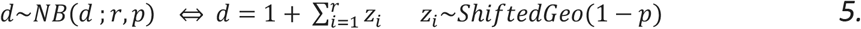

where *ShiftedGeo*(1 – *p*) is *p^z^*(1 – *p*) with z an integer z ≥ 0 [24]. Equality 5 permits to write an HSMM’s transition matrix with NB distributed states, establishing a correspondence to a transition matrix in which each state solely dependent on the previous one (HMM) (figure 1C). The new formulation creates an HMM *embedding* of a HSMM. An HMM embedding permits the use of inference machinery developed for the estimation of parameters in HMMs with a computational complexity that scales as O(r) for each state.

Next, we model the data generating distribution that represents the likelihood that reads in a certain genomic region are generated by open or closed chromatin. The core element of the ATAC-seq assay is a modified version of the Tn5 transposon, binds a 9 bp region when binding to DNA [7]. After preferential binding to open DNA, Tn5 tagments DNA, leaving behind a DNA adaptor. A correctly oriented second event can be used to sequence the intervening fragment to identify tagmented locations [27]. Observations representing Tn5 binding on each individual base pair can be modelled using Bernoulli distribution (Tn5 bound or not bound at a particular base). A complete ATAC-seq assay consists of millions of reads caused by transposition events on thousands of cells, statistically, this is equivalent to repetitive Bernoulli trials. These events can be represented by the one-parameter Binomial distribution (setting as the number of trials the maximum number of binding events on a base pair per chromosome). However, we find that the binomial distribution cannot differentiate between open and closed chromatin effectively due to the sparse nature of each binding event (especially in the case of small sample size and single-cell data sets, supplemental figure 2a). We observed that a geometric distribution better represents the number of events present at each base of open and closed chromatin (supplemental figure 2b), and this completely specifies our initial Bayesian approach. In summary, the presented probabilistic graphical model provides predictive insight into chromatin state and as such defines its accessibility.

**Figure 2:**
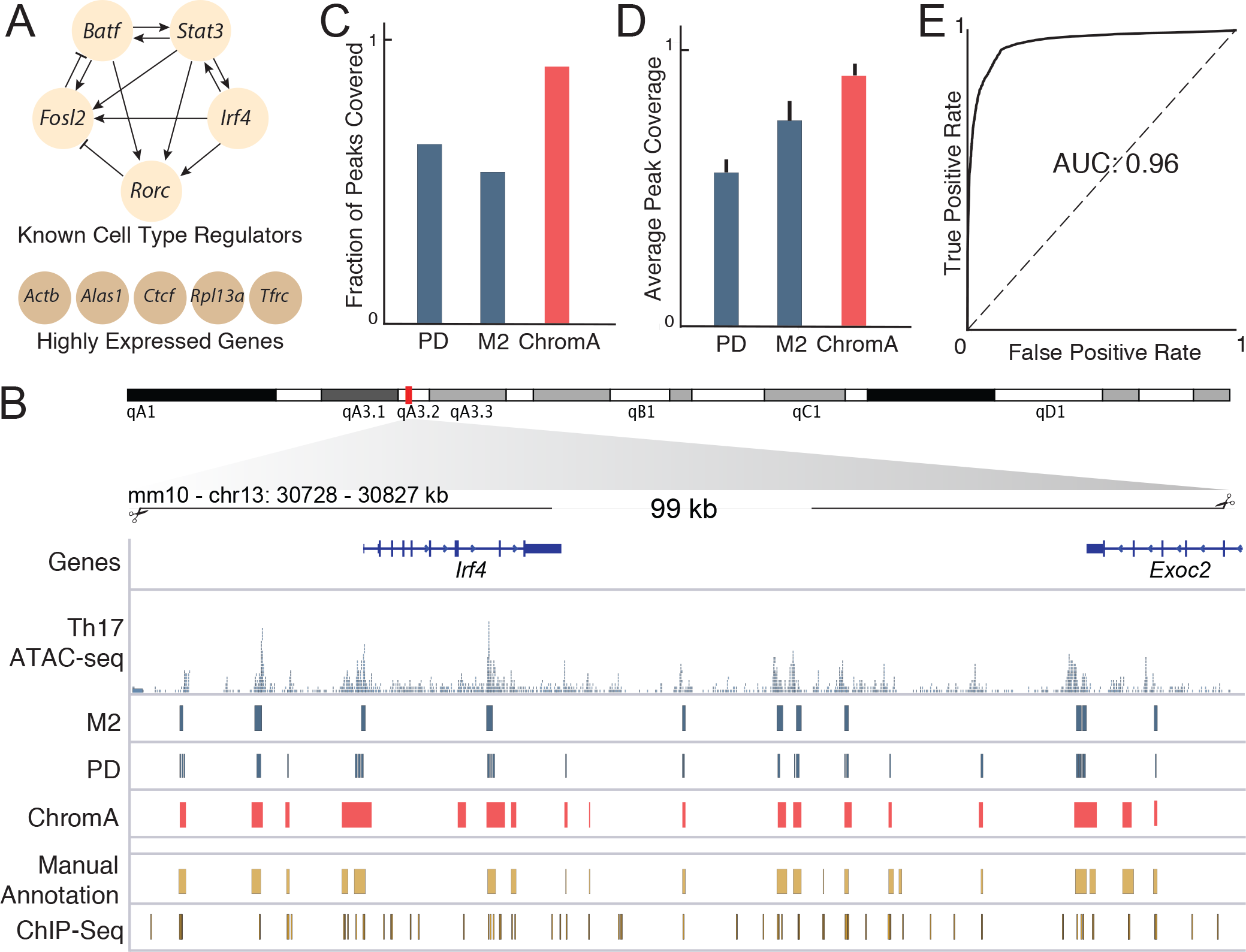
Validation of ChromA on Ground-Truth Datasets. (a) Genomic loci were selected to create a validation data set from highly expressed and transcription factor genes regulating Th17 development. To manually curate genomic regions, information from ChIp-seq and ATAC-seq experiments were combined. (b) Example of curated genomic locus flanking the Actin-b gene in the mouse genome. ATAC-seq, ChIp-seq, peakdeck (PD), macs2 (M2), ChromA and manual annotations are displayed. ChromA algorithm recovers a greater number of ground truth peaks than competing algorithms and covers each peak more thoroughly. (c) Fraction of manually annotated peaks covered with at least one peak. (b) Average fractional area recovered from each peak. (c) Our results are insensitive to the threshold used to annotate chromatin accessibility. ROC curve comparing ChromA against manually ground truth annotations.

### Validating chromatin accessibility annotations

We focused on validating our method on data collected from Th17 cells for which a validated regulatory network delineating their differentiation has been identified [12]. ATAC-seq, several methylation marks, and ChIP-seq on focal transcription factors all of which play a deterministic role in cell fate commitment have been assayed in FACS-sorted Th17 cells [12] [13]. We combine this information to manually annotate chromatin accessibility. We annotate 10 of the best studied loci for this cell-type, each approximately 100kb in size, consisting of regulatory regions surrounding highly expressed genes (Actb, Rpl13a, Alas1, Ctcf, Tfrc) and master-regulator transcription factors (Rorga, Batf, Fosl2, Irf4, Stat3) (figure 2a). We based our curated annotations on the integration of information from three different sources: I) the existence of ATAC-seq regions with higher number of binding events than the surrounding background, ii) the occurrence of H3k27 acetylation marks [11], and iii) the presence of an accumulation of ChIP-seq binding events for the different transcription factors assayed (supplementary figure 3). These annotations serve as ground truth values for comparison, in order to evaluate our model’s performance.

**Figure 3:**
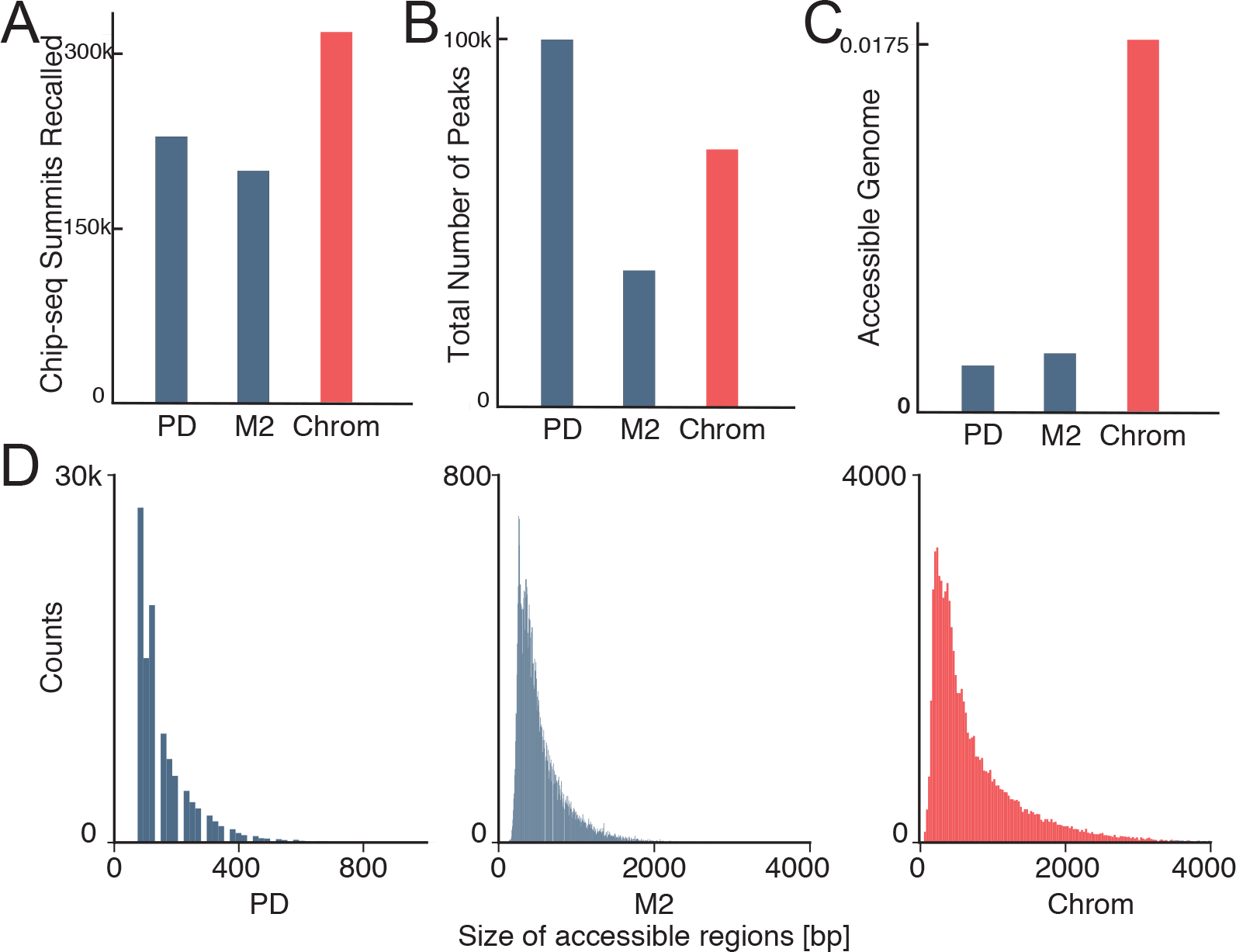
Genome-wide Validation of Chromatin Accessibility Annotations. ChromA outperforms different chromatin accessibility algorithms genome-wide. (a-c) ChromA effective genome-wide performance recalls the highest number of ChIp-seq calls mantaining a reasonable number of peaks. (a) Number of ChIp-seq peaks recalled. (b) Total number of peaks and (c) fraction of the genome annotated as accessible. (d) Distribution of accessible regions for different chromatin annotation algorithms. Histograms depicting size of accessible chromatin regions annotated by peakdeck, macs2 and ChromA. Posterior size distribution of ChromA’s accessible regions resembles a negative binomial distribution, also observed in macs2. peakdeck distribution is highly dependent on algorithmic parameters, such as window size, etc.

Next, to assay chromatin annotations (Figure 2b), we used two different metrics: the fraction of the total number of manually annotated peaks that contains at least one peak generated using ChromA (Fraction of peaks covered), and the average fraction of coverage of each peak (average peak coverage). We used these metrics to compare ChromA annotations against peakdeck [20] and macs2 [29], two of the most commonly used tools to annotate ATAC-seq data. In both cases, ChromA annotations not only recovered a higher fraction of correctly annotated peaks but also on average generated better coverage of each of the accessible regions (figure 2c-d). ChromA annotations are performed by thresholding the inferred posterior estimate of chromatin state. This algorithmic parameter does not play a major factor in ChromA’s annotations, highlighting the robustness of our model (figure 2e, AUC = 0.96).

Next, we examine ChromA’s performance genome-wide, again focusing on Th17 cells. In this case, manual annotation is not feasible for computing a ground truth metric (with changes in chromatin accessibility spanning the full genome [12] [13]). Instead, we reasoned that ChIP-seq locations can be used as a proxy to indicate chromatin accessible regions and therefore used ChIP-seq data for validation experiments. Compared to other existing methods, ChromA’s predictions faithfully recover the greatest number of ChIP-seq calls, while maintaining a comparable total number of peaks. In addition, ChromA annotates the highest genome fraction as accessible, consistent with ChIP-seq information (figure 3a-b-c). While, macs2 and ChromA exhibit NB distributed sizes, Peakdeck exhibit a discontinuous size distribution in which an algorithmic parameter (peak size parameter) is a major determinant of its shape (figure 3d). Lastly, we validated ChromA’s performance on four additional data sets, two of them consisting of Th17 cells and the remaining two consisting of CD4+ cells differentiated into Th17 cells (Table 1). To differentiate CD4+ into Th17, CD4+ sorted cells were purified by cell sorting and cultured for 48hrs in Th17 differentiating media [12]. On these data sets, ChromA’s recovered on average 45% more peaks than macs2 (supplementary figure 4). Taken together, these results established ChromA as a top performing tool for discovering accessible chromatin regions from ATAC-seq datasets.

**Figure 4:**
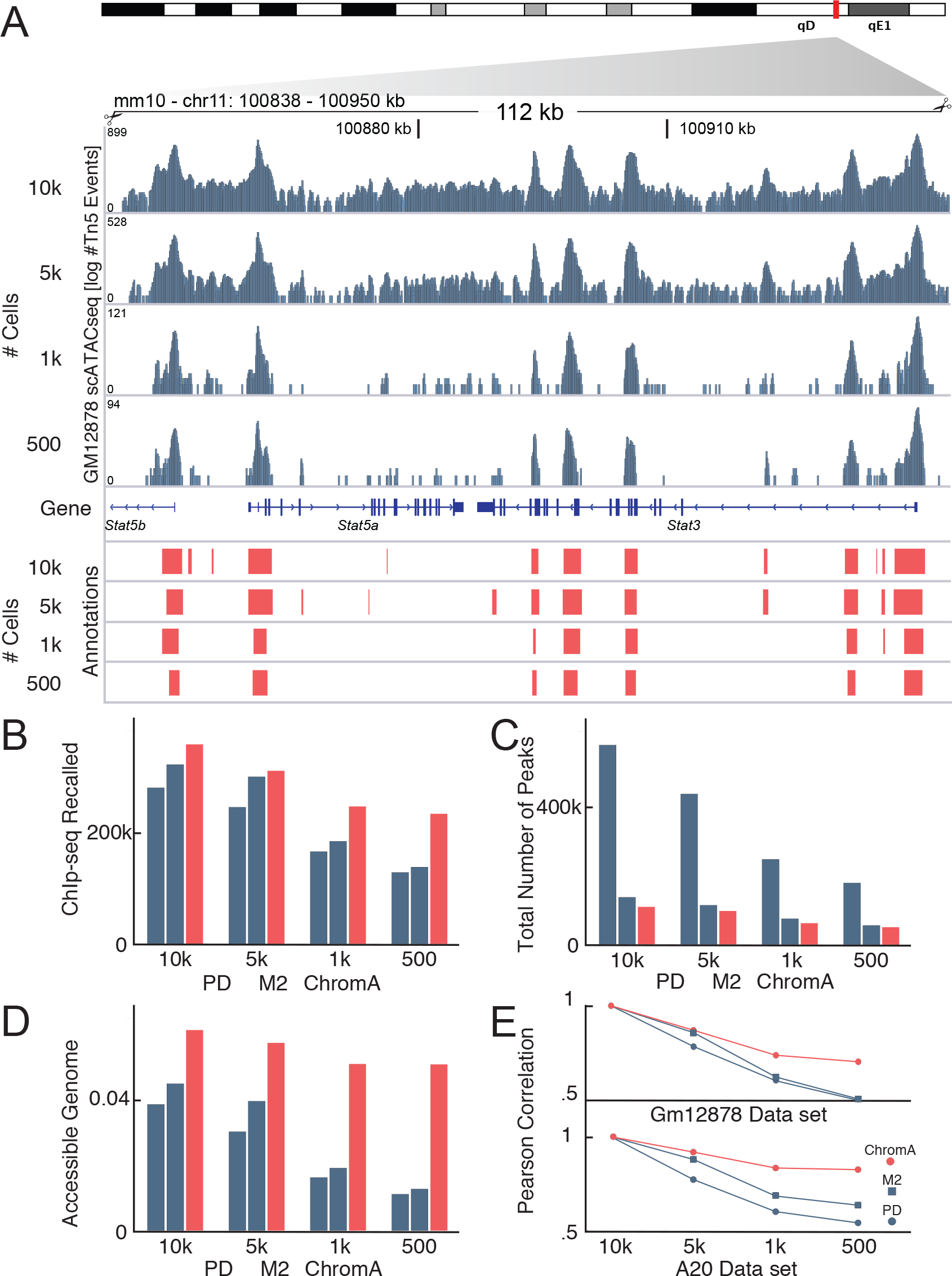
ChromA Annotations Generalize to Single-cell Datasets. (a) Annotations of mouse A20 single cell datasets at the Stat3 genomic locus. Cells are down-sampled from 10000 to 500 cells. ChromA annotations are consistent at different cell depths. (b-e) ChromA extends its effective genome-wide performance to single-cell data sets, again, recalling the highest number of ChIp-seq calls in GM12878 single-cell data sets. ChromA is particularly effective at low cell depths. (b) Number of ChIp-seq peaks recalled, (c) total number of peaks and (d) fraction of the genome annotated as accessible for each down sampled data-set. (e) Correlation between annotations at different cell depths calculated against the entire dataset possessing 10000 cells.

### Chromatin accessibility annotations from single-cell measurements

With minor modifications we can extend ChromA’s core model beyond bulk processing to characterize chromatin accessibility at the single cell level (we describe modifications to the model for hybrid experimental designs that include bulk and single cell data in the next section). Here, we focus our analysis on single cell data sets of mouse B lymphocyte A20 and human lymphoblastoid GM12878 cells (dataset obtained from 10x Genomics, see methods section for a description of the samples). Single-cell datasets exhibit higher dynamic range (DR) than their bulk counterparts (bulk DR ∼ 4 bits, single-cell DR ∼ 11 bits; Supplemental Figure 5). To characterize single-cell chromatin datasets and compare them to bulk datasets, we employ a set of metrics aimed to quantify dataset quality. We compute a signal to noise ratio (SNR) centered around gene promoter regions, the fraction of reads in accessible regions, and a ratio between read lengths centered around mono-nucleosome and nucleosome-free regions (supplemental methods, Table 1, supplemental Figure 6).

**Figure 5:**
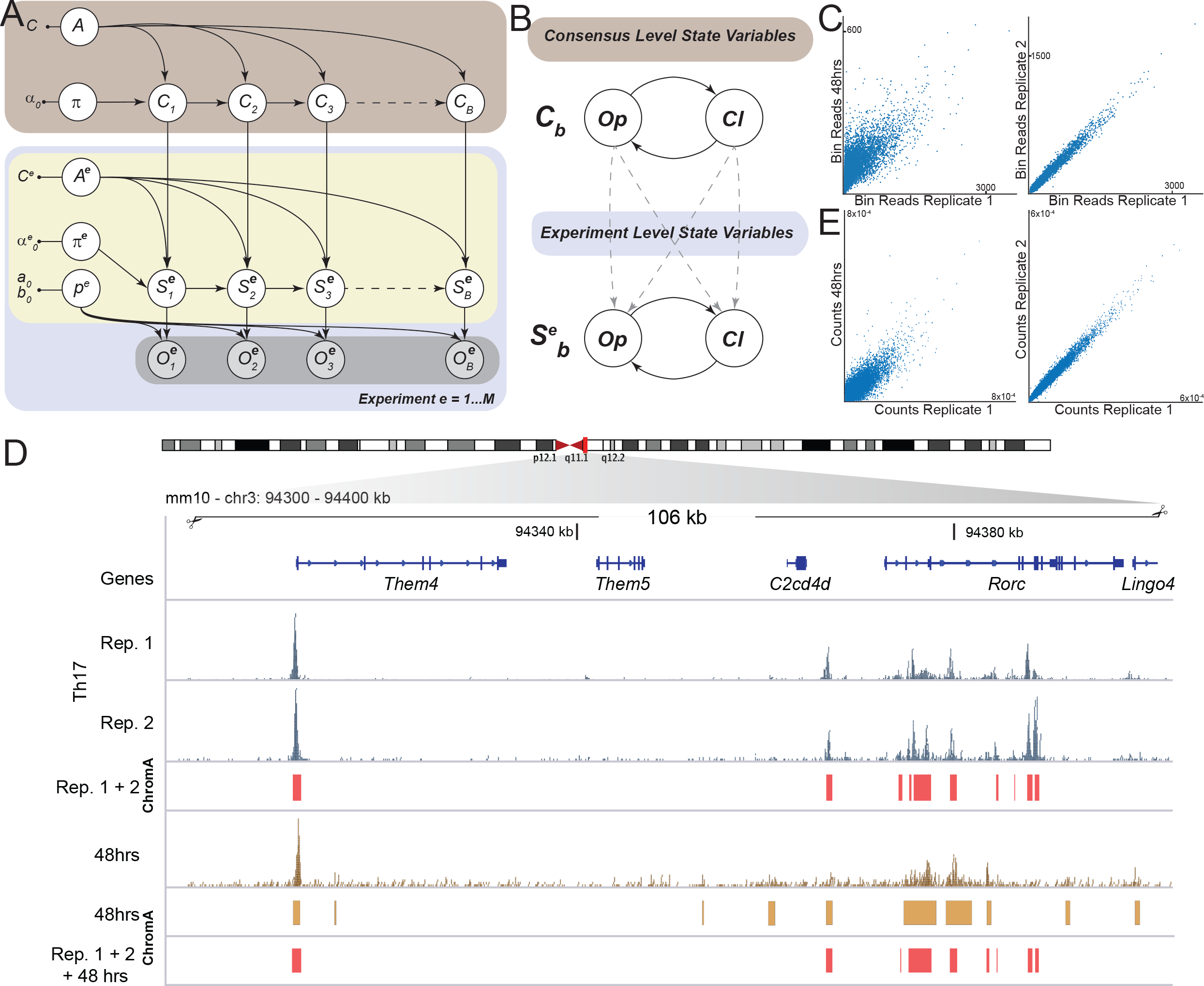
Consensus ChromA Integrates Information from Different Data Sets and Replicates. (a) Consensus ChromA probabilistic hierarchical graphical model. The model is divided into a top (light brown) and a bottom layer (light blue). The bottom layer shows ChromA’s probabilistic model for each data set analyzed (akin to figure 1.b). The top layer schematic shows how consensus variables, *C*, are explicitly linked to latent state variables for each replicate (or integrated experiment) according to Markovian dynamics. *π*, *p_g_* and *A* variables are as in figure 1.b. (b) Consensus and experiment variables *C* and *S^e^* evolve alternating between open, Op, and closed, Cl, states. The top and the bottom layers are linked by introducing into each experiment a dependency on the state of the consensus variable. This dependency is represented in a transition matrix h. Prior information on the parameters of the matrix encode our belief that at each base, experimental states should be more likely to transition into the state of the consensus variable. (c) Raw read correlation between replicates of sorted Th17 cells’ datasets (right) and Th17 cells against CD4+ cells incubated in Th17 differentiation media for 48hrs (left). (d) Consensus ChromA annotations integrates information from different replicates creating and deleting accessible regions based on context. Raw reads from sorted Th17 cells’ replicates and sorted CD4+ cells at a genomic locus. ChromA annotations for single CD4+ dataset. Consensus ChromA annotations for Th17 replicates and Th17 replicates and CD4+ cells are shown in red. (e) ChromA Consensus creates a common representation of chromatin accessible regions. When both Th17 replicates are combined together with CD4+ cells, the resulting consensus representation maintains the high correlation observed only when both Th17 cells’ replicates are used. CD4+ peaks are filtered, and only correlated peaks survived.

To study the robustness of ChromA’s single-cell approach, we varied the total number of cells in our data sets and studied how chromatin annotations varied as we down-sampled data to different depths. Next, we annotate human and mouse single cell datasets with ChromA and competing approaches (Figure4a) and assess the accuracy of genome wide annotations using ChIP-seq locations, available solely for GM12878 cells (supplementary methods). Once more, ChromA recovered the highest number of ChIP-seq calls and annotated the highest accessible genome fraction at every cell depth, consistent with ChIP-seq information (Figure 4b-c-d). ChromA features generalize at the single cell level, allowing the recovery of broad peaks, which are ideal for transcription factor foot-printing. Thus, our explicit modeling of length/duration via HSSM makes ChromA particularly robust to under-sampling. When cells are down-sampled and data is sparse, the correct treatment of regulatory element’s durations by ChromA’s HSMM permits correct annotation of accessible chromatin regions. By contrast, other methods tend to fragment annotations into many disjointed loci or select the null model inappropriately (Figure 4b-e).

For both bulk and single-cell data sets, ChromA uses approximate Bayesian algorithms to perform scalable inference. Moreover, additional acceleration is achieved through biologically inspired approximations (Supplementary Section 1, Supplementary Figure 7-8). Taken together, our computational experiments validate our algorithms as an effective platform for chromatin annotation under different experimental settings.

### Integrating Information from Different Replicates to Create a Consensus Chromatin Annotation

High throughput genomic data sets can suffer from high variability, making experimental replication an essential element of any genomics experimental design. Bayesian inference is ideally suited to accommodate many-trial experiments. We design ChromA to infer a consensus chromatin state representation by harnessing the statistical power from different experimental replicates, thus inferring a more confident posterior estimate. Furthermore, a consensus representation can gather information from both single-cell and bulk experiments, combining both platforms to inform chromatin state. In addition, when analyzing experiments from different cellular populations, this mode facilitates both the creation of a shared peak universe and the identification of sample-specific peaks. Therefore, consensus ChromA probabilistic model enables the processing of *replication*, the *integration* of different experimental platforms and, the *comparison* of diverse cellular populations.

Our model consists of consensus and individual experiment chromatin state variables (indicated with letter *C* and *S^e^*, respectively; figure 5a). The challenge in this context is to integrate state persistence into each experimental state sequence, while augmenting the model to share information among experiments (while maintaining computational tractability and scalability). To execute these goals coherently, we maintain negative binomial HSMM dependencies in our consensus chromatin state variables, *C*. Next, variables *S^e^* behave under semi-Markovian dynamics and incorporate a dependency on the state of the consensus representation. To model this dependency, we resort to the HMM NB-embedding of the HSMM. We augment individual experiment NB-embedding to include a transition matrix depending on the consensus representation (supplemental equations, figure 5). The link between each experiment and the consensus representation is possible because the HMM NB-embedding, indicated with C, S^e^ below, creates a base by base dependency. This dependency can be tailored to connect individual replicates or data-sets (and their embeddings, S) to our consensus model, C, as follows. We begin by writing the probability that represents the transition matrix in this model.

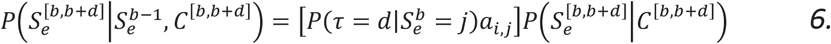

where the letter *e* is an index for each experimental replicate. Equation 6 represents the HSMM probability of transitioning from a state at base b-1 into a state spanning bases b to b + d, given consensus variables at those bases. This probability factorizes into a HSMM transition term times a term linking each experiment to the consensus variables. We rewrite the previous equality by using the HMM NB-embedding transition matrix, *A^e^*, and a base-by-base consensus link transition matrix *H*.

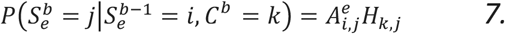

Thus, the segmentation of each experimental data set should be made based on the observed density of the states throughout the entire sequence. Finally, we use prior information to encode our belief that each experimental sequence is more likely to transition into an open state if the current consensus state is open rather than closed. We named our consensus approach consensus-ChromA.

To demonstrate the model’s efficacy in integrating information from replicate experiments, we apply this new statistical tool to an ATAC-seq data set comprised of two replicates of Th17 sorted cells. The raw signals of replicates are highly correlated (correlation coefficient = 0.99), measured by correlating binned raw bam files using deeptools ([31], multiBamSummary). Additionally, to study the model’s robustness to outliers, we select a dataset of our naïve CD4+ T cells cultured in Th17 conditions for 48 hours. This dataset of differentiated Th17 cells exhibits lower correlation when compared against Th17 cells (correlation coefficient = 0.68, figure 5c). How does read correlation translate into chromatin accessibility correlation? To answer this question, we segment accessible regions using consensus-ChromA in two steps: first, we segment both Th17 replicates; second, we annotate both replicates together with 48hrs cultured cells (figure 3d). Bases in which mismatches occurred experience a trajectory consistent with the approximate finite memory of the process by pooling information from every replicate (supplemental figure 9). Bases in which every replicate agrees, generate a consensus state coinciding with the mutual state of every replicate.

To assess our model’s performance, we measured the level of correlation among data sets based on the number of Tn5 transposition events occurring at each accessible chromatin region. In this case, replicates continue to be highly correlated, as expected (correlation coefficient = 0.99, accessible regions calculated with consensus-ChromA run only on Th17 cells replicate 1 and 2). This correlation remains unaltered even, when the outlier is included into the analysis (correlation coefficient = 0.99, accessible regions calculated with consensus-ChromA run on Th17 cells replicate 1, 2, and 48hrs cultured CD4+. figure 4e). Although consensus-ChromA builds accessible regions common to the three data sets, this common basis does not alter the fact that 48hrs cultured CD4+ cells correctly stand as an outlier, the individual model, S, for the 48hrs cultured CD4+ cells is not perturbed (correlation coefficient Replicate 1 vs 48hrs = 0.651, correlation coefficient Replicate 2 vs 48hrs = 0.655; accessible regions calculated with consensus-ChromA run on Th17 cells replicate 1, 2, and 48hrs cultured CD4+. figure 3e).

## Discussion

A major goal in epigenomic analysis is to systematically dissect chromatin accessible regions in cell types under different conditions. We developed ChromA, a powerful probabilistic model for the analysis of ATAC-seq experiments as a means to annotate chromatin accessible areas in the genome. We validated our approach with curated regions in the mouse genome and by assessing our algorithm performance against chromatin immunoprecipitation binding events, a proxy of accessible chromatin. The correlation between binding events and the ATAC-seq signal permitted us to directly assess accessibility. To test the generalizability of our method, we tested our algorithm in single cells. Here, we demonstrate that our probabilistic algorithm is useful both in single cell and aggregate populations. These analyses show that our method can be readily extended to more complex models as new technologies and more complex experimental designs emerge.

We found that ChromA facilitates complete genome chromatin accessibility annotations for human and mouse genomes from ATAC-seq information. Our algorithm has several advantages over previous approaches. First, ChromA recovers wider accessible regions, facilitating transcription factor foot-printing. ChromA also exhibits higher sensitivity allowing for the recovery of less prominent peaks. As a result, single cells datasets, in which there is an extended dynamic range compared to bulk measurements can also be analyzed with our software. Finally, by integrating different experiments, ChromA is able to create a consensus annotation and thereby increase the signal-to-noise. This consensus representation is not a simple intersection or sumation of the experiments under analysis. Rather, ChromA combines information by merging and intersecting accessible regions of individual experiments, as well as including new accessible regions, as information and context demands. This analysis indicates that additional insights can be extracted by integrating different sources of information. In the future, we plan to extend ChromA to integrate different experimental procedures, extracting and combining information from a wide range of approaches [32, 33]. Given the broad importance of ATAC-seq information in mapping the chromatin landscape and the advent of commercial platforms to reliably expand the technique at the single-cell experimental designs, ChromA provides a useful tool to analyze such data and generate unique insights into epigenetic regulation.

In summary, our hierarchical probabilistic model reaches a more robust (in a formal statistical sense) posterior estimate of chromatin state than its single experiment counterparts by sharing information across replicates. This demonstrates the flexibility of our generative model, which can be further elaborated to incorporate complex dependencies between chromatin states. Finally, our consensus annotation method provides a principled manner of integrating bulk and single cell experiments to create robust annotations.

## Online Methods

### Bulk ATAC-seq libraries and Pre-processing

ATAC-seq libraries were downloaded from NCBI’s GEO Database under accession GSE113721. The following preprocessing pipeline was used to generate aligned reads. Adapters were trimmed using cutadapt. Reads were aligned using BOWTIE2 to the murine mm10 reference genome and then filtered for mapping quality greater than Q30. Duplicates were removed using PICARD (http://picard.sourceforge.net) and subsequently, mitochondrial, unmapped and chromosome Y reads were removed. For peak-calling, ChromA corrects the read start sites to represent the center of the tagmentation binding event, the + strand were offset by +4 bp, and all reads aligning to the – strand were offset −5 bp. Additionally, ChromA filters peaks using a custom list that combines blacklisted genomic regions from the ENCODE project (http://mitra.stanford.edu/kundaje/akundaje/release/blacklists/mm10-mouse/mm10.blacklist.bed.gz). This filtering step takes place when building the set of transposition events by removing all the events falling into the blacklisted regions.

### Single-cell ATAC-seq libraries

Single cell datasets were downloaded from 10X genomics https://support.10xgenomics.com/single-cell-atac/datasets/1.0.0/atac__hgmm_10k. Briefly, they consist of a mixture of fresh frozen human (GM12878) and mouse (A20) cells collected with the Chromium Single Cell ATAC platform, and demultiplexed and pre-processed with the single-cell ATAC Cell Ranger platform. Cells were sequenced on Illumina NovaSeq with approximately 42k read pairs per cell. Down-sampled datasets are provided from the online website. Tsv files are provided listing binding events. ChromA incorporates the ability of importing tsv files directly from Cell Ranger pipelines.

### Dataset Metrics

ChromA reports different quality control metrics to assess dataset quality. Given ChromA annotations, the fraction of reads in peaks (FRIP) is calculated as the number of reads laying within peaks versus the total number of reads in chromosome 1. This is calculated using properly paired and mated reads. Signal to noise ratio (SNR) is calculated by defining promoter regions in the mouse or human genome as regions spanning 1kb upstream, 3kb downstream from gene start sites. Insert size distribution is reported as an additional file and insert size metric is computed as the ratio between the number of reads with insert size between 190-210bp to the number of reads with insert size between 60-80bp for chromosome 1. Finally, we extrapolate the number of properly paired and mated reads by computing that number for chromosome 1 and multiplying by the total length of the genome and then dividing by the length of chromosome 1.

### Detection of Chromatin Accessible Regions

To perform experiments to validate our algorithm, we ran ChromA in each sample individually using standard priors (described below). An example of running ChromA on a wild type dataset of Th17 cells, using the mouse genome with our bulk model is detailed next: chrom.py -b “Th17_1_noMito.bam" --species mouse -sp th17_wt1.pkl -sb th17_wt1.bed -inf mo -sen low >> logwt1l.log. We ran peakdeck (parameters -bin 75, -STEP 25, -back 10000, -npBack100000). Peaks were identified using the macs2 software. We run macs2 using two sets of parameter and always compare against the best performing set (parameters: -m 10,30 -g 1865500000 --bw=200 or --nomodel --shift -100--extsize 200 --broad --keep-dup all).

### Transcription Factor Binding Prediction

TF ChIP-seq and control sequencing data were downloaded from GEO (GSE40918), mapped to the murine genome (mm10) with bowtie2 (2.2.3), filtered based on mapping score (MAPQ > 30, Samtools (0.1.19)), and duplicates removed (Picard). Peaks were identified using the macs2 software (version 1.4.2) using the settings (parameters: -m 10,30 -g 1865500000 --bw=200) and retained for raw p-value < 10-10. All data sets were processed against an appropriate control. We retained summit locations to create a binding event localizing at a particular base pair.

Transcription factor binding events for Gm12878 were downloaded from ReMap [31] (http://pedagogix-tagc.univ-mrs.fr/remap/celltype.php?CT=gm12878) by filtering the database for the cell type gm12878. There are 131 TFs in this database that correspond to the particular cell line, among which we can find CTCF, PouxFx factors and members of the Pax, Stat and Etv families.

### Validation of ChromA annotations

To compare ChromA against different algorithms we used different metrics, the fraction of peaks covered, average peak coverage and total coverage. We compute each metric from the intersection of bed files originating from the manually annotated regions versus algorithmically annotated regions. To compute the fraction of peaks covered (fpc), each manually annotated peak is intersected with the list of peaks algorithmically generated. If the intersection returns non-empty bases, the peak is considered intersected and recorded as such. The final metric value is computed by dividing the number of intersected peaks over the number of peaks 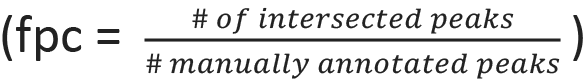 To compute the average peak coverage (apc), we again intersect each manually annotated peak and count the base pairs in the intersection over the total number of base pairs in the peak 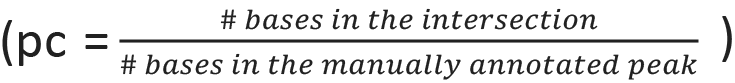 The apc is computed as the mean of the pc for every manually annotated peak. We report the apc as mean +/− sem. To compute the total coverage (tc), we add all the intersected bases and divide by the total number of bases in manually annotated peaks 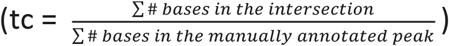.

### ChromA model and core Algorithm

ChromA’s probabilistic graphical model is described throughout the text. Here, we present in more detail the entire generative process. The observed number of Tn5 binding events X_b_ at each base b is drawn independently through the process here described.

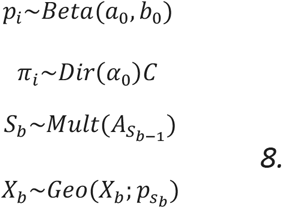

S_b_ denotes the chromatin state at base b, and it is distributed according to the transition matrix at the previous state. p is the probability of observing X_b_ number of binding events at base b given the current chromatin state. Z_j_, k_j_, n_j_ are prior parameters. A denotes the HMM embedding of the HSMM and for a two-states model with 3 and 2 states it is written as:

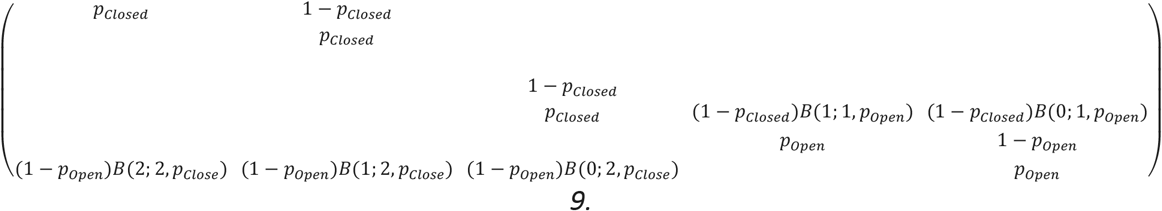

we used 5 and 2 as our fixed number of states and although we perform computational experiments to fit p, these values were fixed at 1_x_10^−4^. a_0_ and b_0_ parametrize pseudo-counts for the probability of observing a number of binding events in a particular base. We set these values to (1, 50) for the state that represents closed chromatin and (20, 10) for the state that represents open however, results are insensitive to these values. *α*_0_ denotes the prior pseudo-counts for the initial state of the Markov process. Given our strategy that identifies batches surrounded by empty regions, we assume that the process starts in the closed state, *α*_0_ = (1000, 1). Again, the algorithm is insensitive to this value, as only the first few bases will be affected by it. Pseudo-code of ChromA single and multiple experiments annotations is described in supplemental figure 8.

**Table 1: Quality Metrics for Bulk and Single-Cell ATAC-seq Data Sets.**

## Software availability

A Python implementation of ChromA is available for download on GitHub: https://github.com/marianogabitto/ChromA The website will be updated periodically with new versions. To install, run: pip install git+ https://github.com/marianogabitto/ChromA.

## Data availability

The accession number for the single-cell and bulk ATAC-seq data reported in this paper is GEO:. [[and repeat citations, does 10x-scATAC have a geo ?]]

## Acknowledgements

This research was supported by the Simons Foundation. We would like to thank P. Shah, B. Olsen and G. Zheng from 10x Genomics for initial discussions and comments on the manuscript. We would like to thank M. Teper, E. Macosko, L. Paninski and E, Mazzoni for their critical comments on the manuscript.

## Supplementary Figure Legend

**Supplementary Figure 1: The Negative Binomial Distribution Effectively Describes Genomic Elements.** (a) Example of negative binomial distribution with parameters r=3, p=0.1 and geometric distribution with parameter p=0.1. The mode of the geometric distribution is always 0, which is not ideal to model genomic elements. (b) Gene length is correctly described by a negative binomial distribution. Histogram of gene sizes in chromosome 4 of mouse. Mouse genome annotation was downloaded from Ensembl, https://useast.ensembl.org/Mus_musculus/Info/Annotation.

**Supplementary Figure 2: The Number of Tn5 Binding Events at Open and Closed Chromatin can be Described by a Geometric Distribution.** (a) Histogram depicting the number of Tn5 binding events in manually annotated open and closed chromatin. Both distributions possess their maximum at 0 and can be effectively described with a geometric distribution. (b) Parameter posterior estimates calculated for a Bernoulli, binomial or geometric distribution using the number of Tn5 binding events in open and closed chromatin. Each distribution possesses a random probability measure, p. The gap between the mean open and close chromatin measures, p, is computed as the difference between the mean value of p at each state.

**Supplementary Figure 3: Example Manual Annotations on Selected Genomic Regions.**Manual annotations on two out of ten selected genomic regions. ChIp-seq binding locations collected for different transcription factors are used to delineated chromatin accessible regions.

**Supplementary Figure 4: Extended Validation of ChromA Annotation Algorithm.** (a, b, c) We validate ChromA’s performance by annotating the same genomic loci in 5 datasets of wild type and CD4+ treated after 48hr Th17 cells and re-computing validation metrics.

(a) Average fraction of manually annotated peaks covered with at least one peak.

(b)Average fractional area in which a peak is covered.

(c)Average total area covered on our manual annotations.

(f-h) Entire genome annotations.

(f) Number of ChIpseq peaks recalled, (g) total number of peaks and (h) fraction of the genome annotated as accessible.

**Supplementary Figure 5: Single-cell Data Sets Exhibit Extended Dynamic Range Compared to Bulk data sets.** Histogram of Tn5 binding events in single-cell data set of 10000 gb12878 human cells (left) and bulk Th17 mouse cells (right). Counts are depicted in log scale. Due to high level of sparsity, the number of reads in bulk dataset do not typically surpasses more than 50 Tn5 binding events per base. This is in contrast to single cell datasets in which the number of Tn5 binding events can reach on the order of thousands.

**Supplementary Figure 6: Insert Size Distribution for bulk and Single Cell Datasets.** Histogram depicting insert size distribution computed for every dataset used in this work. In every plot, first peak corresponds to nucleosome-free regions and subsequent peaks correspond to mono, bi and tri nucleosome reads.

**Supplementary Figure 7: Scaling of ChromA Parameters’ Inference.** Parameter inference in ChromA (performed for each batch of data or set of replicate datasets) is accelerated by distributing computational load in batches.

(a) Chromosomal regions are divided into batches of data by identifying flanking regions lacking a significant number of reads. These batches of data have a minimum length of 100 kbp and experimentally determine maximum of less than 600kbp.

(b) To ensure computational scalability, we compare parameter inference with no batch acceleration (Full) against stochastic (SO) and memoized optimization (MO). Algorithms run for 10 iterations on a dataset composed of chromosome 19 of Th17 cells using our two-state bulk inference algorithm. MO and SO algorithms converge faster when compared to the Full algorithm, seeing as a faster convergence of the expected lower bound and the open and closed chromatin states’ parameters.

(c) Entire chromosomes can be annotated by using open and closed states’ parameters that generalize chromosome wide. Histogram depicting open and closed probability parameters fit independently in each data batch. On red is the posterior parameter value when using a single set of open and closed parameters chromosome wide.

**Supplementary Figure 8: ChromA Running Time and Complete Computational Pipeline.**

(a) ChromA running time of one iteration on mouse chromosome 1 (mean +/− sem). Minimal computational overhead is observed when using batch algorithms. Full refers to our algorithm that uses no batches. MO: memoized optimization and SO: stochastic optimization.

(b) ChromA computational pipeline.

**Supplementary Figure 9: Consensus Annotations Integrate Context Information from Different Data Sets.** (a, b, c, d) Chromatin annotations are integrated using two replicates of wild type Th17 sorted cells. Many examples throughout the genome highlight that annotations are not just intersection or union of replicates’ annotations.

(a) Extracted genomic region as an example to highlight intersection of replicates in region 1 (b), rescue of lost peak in region 2 (c), and addition of a missing peak in region 3 (d).

(b) In region 1, chromatin annotations resemble a simple intersection of replicates’ annotations.

(c) In region 2, a peak only present in one replicate is annotated on the consensus representation, presumably, because of the presence of an accumulation of reads in both data sets.

(d) In region 3, no replicate includes a peak in the genomic locus however, when information is combined, the consensus state discovers missing information.

(e)-(f) Example of chromatin consensus annotation in which we include posterior chromatin consensus state variable before thresholding.

## References

1. Kornberg R.D. Chromatin structure: a repeating unit of histones and DNA. Science. 1974 May 24; 184(4139):868–71.

2. Kornberg RD, Lorch Y. Chromatin structure and transcription. Annu Rev Cell Biol. 1992;8:563–87.

3. Mellor J. The dynamics of chromatin remodeling at promoters. Mol. Cell. 2005, 19(2):147–57.

4. Mitchell P.J., Tjian R. Transcriptional regulation in mammalian cells by sequence-specific DNA binding proteins. Science 1989; 245: 371–378.

5. Kohwi M and Doe C. Q. Temporal fate specification and neural progenitor competence during development. Nature Reviews Neuroscience. 2013. Volume 14, pages 823–838.

6. Slattery M., Zhou T., Yang L., Dantas Machado A.C., Gordân R., Rohs R. Absence of a simple code: how transcription factors read the genome. Trends Biochem Sci. 2014.39(9):381–99.

7. Zhou L., Littman DR. Transcriptional regulatory networks in Th17 cell differentiation. Curr Opin Immunol. 2009 Apr;21(2):146–52.

8. Systems biology. Conditional density-based analysis of T cell signaling in single-cell data. Krishnaswamy S1, Spitzer MH2, Mingueneau M3, Bendall SC2, Litvin O1, Stone E4, Pe’er D5, Nolan GP2.

9. Papalexi E, Satija R. Single-cell RNA sequencing to explore immune cell heterogeneity. Nature Reviews Immunology. 2017.

10. Stubbington MJT, Rozenblatt-Rosen O, Regev A, Teichmann SA. (2017) Single-cell transcriptomics to explore the immune system in health and disease. Science Oct 6; 358(6359):58–63

11. Histone H3K27ac separates active from poised enhancers and predicts developmental state. Menno P. Creyghton, Albert W. Cheng, G. Grant Welstead, Tristan Kooistra, Bryce W. Carey, Eveline J. Steine, Jacob Hanna, Michael A. Lodato, Garrett M. Frampton, Phillip A. Sharp, Laurie A. Boyer, Richard A. Young, and Rudolf Jaenisch. PNAS December 14, 2010 107 (50) 21931–21936.

12. Ciofani M, et al. A validated regulatory network for Th17 cell specification. Cell. 2012 Oct 12;151(2):289–303.

13. Miraldi E. R., et al. Leveraging chromatin accessibility for transcriptional regulatory network inference in T Helper 17 Cells. BioArxiv. doi: http://dx.doi.org/10.1101/292987.

14. ATAC-seq: A Method for Assaying Chromatin Accessibility Genome-Wide. Jason Buenrostro, Beijing Wu,Howard Chang, and William Greenleaf. Curr Protoc Mol Biol. 2015; 109: 21.29.1–21.29.9.

15. Buenrostro JD, et al. Single-cell chromatin accessibility reveals principles of regulatory variation. Nature. 2015 Jul 23;523(7561):486–90.

16. Inoue F, Kircher M, Martin B, et al. A systematic comparison reveals substantial differences in chromosomal versus episomal encoding of enhancer activity. Genome Res. 2017;27(1):38–52. PMID: 27831498; PMCID: PMC5204343

17. Lizio M, et al. Update of the FANTOM web resource: high resolution transcriptome of diverse cell types in mammals. Nucleic Acids Res 45: D737–D743 (2017). 10.1093/nar/gkw995.

18. Macosko E*, Basu A, Satija R, Nemesh J, Shekhar K, Goldman M, Tirosh I, Bialas, A, Kamitaki N, Martersteck E, Trombetta J, Weitz D, Sanes J, Shalek A, Regev A, McCarroll S*. “Highly Parallel Genome-Wide Expression Profiling of Individual cells Using Nanoliter Droplets.” Cell 2015, 161, 1202–1214.

19. The accessible chromatin landscape of the human genome. Nature489:75–82, 2012.

20. Cusanovich D. A. et al. The cisregulatory dynamics of embryonic development at single cell resolution. Nature 555, 538–542 (2018).

21. Daugherty A. C. et al. Chromatin accessibility dynamics reveal novel functional enhancers in C. elegans. Genome Res. 27, 2096–2107 (2017).

22. Jorja G. Henikoff, Jason A. Belsky, Kristina Krassovsky, David M. MacAlpine, and Steven Henikoff. Epigenome characterization at single base-pair resolution. PNAS November 8, 2011 108 (45) 18318–18323.

23. Boyle, AP; Davis S; Shulha HP; Meltzer P; Margulies EH; Weng Z; Furey TS; Crawford GE (2008). “High-resolution mapping and characterization of open chromatin across the genome”. Cell. 132 (2): 311–22.

24. Johnson M. J. and Willsky A. S. Stochastic Variational Inference for Bayesian Time Series Models. ICML-14. 2014. 1854–1862.

25. Durbin R., Eddy S. R., Krogh A., Mitchison G. Biological Sequence Analysis: Probabilistic Models of Proteins and Nucleic Acids 1st Edition.

26. Guédon Y. Estimating Hidden Semi-Markov Chains from Discrete Sequences. Journal of Computational and Graphical Statistics. 2003. Vol. 12, 3. pp. 604–639.

27. Adey A., Morrison H.G., Asan, X. X, Kitzman J.O., Turner EH, Stackhouse B, MacKenzie AP, Caruccio NC, Zhang X, Shendure J. Rapid, low-input, low-bias construction of shotgun fragment libraries by high-density in vitro transposition. Genome Biol. 2010; 11(12):119.

28. McCarthy M. T. and O’Callaghan C, A. PeaKDEck: a kernel density estimator-based peak calling program for DNaseI-seq data. Bioinformatics. 2014 May 1; 30(9): 1302–1304.

29. Feng J., Liu T., Qin B., Zhang Y., Liu X. S. Identifying ChIP-seq enrichment using MACS. Nat Protoc. 2012 Sep; 7(9).

30. Ramirez, F., et al. deepTools2: A next Generation Web Server for Deep-Sequencing Data Analysis. Nucleic Acids Research, 2016.

31. Cheneby J., Gheorghe M., Artufel M., Mathelier A., Ballester, B. ReMap 2018: An updated regulatory regions atlas from an integrative analysis of DNA-binding ChIP-seq experiments. Nucleic Acids Research, gkx1092.

32. Rao S. S. P., Huntley M. H., Durand N. C., Stamenova E. K., Bochkov I. D., Robinson J. T., Sanborn A., Machol I., Omer A. D., Lander E. S. and Lieberman Aiden E. A three-dimensional map of the human genome at kilobase resolution reveals principles of chromatin looping. Cell. 2014 Dec 18; 159(7): 1665–1680.

33. Dostie J, Richmond TA, Arnaout RA, Selzer RR, Lee WL, Honan TA, Rubio ED, Krumm A, Lamb J, Nusbaum C, Green RD, Dekker J. Chromosome Conformation Capture Carbon Copy (5C): a massively parallel solution for mapping interactions between genomic elements. Genome Res. 2006 Oct; 16(10):1299–309.

